# Mechanosensitive Remodeling Sustains Rigidity Homeostasis and Elastic Memory in Actin Cortex Models

**DOI:** 10.1101/2025.10.05.680567

**Authors:** Haina Wang, Marco A. Galvani Cunha, John C. Crocker, Andrea J. Liu

## Abstract

The actin cortex is a dynamic biopolymer network that, despite constant architectural changes through the assembly and disassembly of filaments and crosslinkers, sustains a rigid homeostatic state, i.e., a steady state of positive elastic moduli, and exhibits an elastic memory that far outlasts the structural turnover time scale. We still lack simple models that maintain collective rigidity and elastic memory at long times compared to the turnover time. To address this, we develop two complementary elastic network models in which rigidity homeostasis and long-lived elastic memory with complete turnover emerge as a result of mechanosensitive dynamics of filaments (edges) and crosslinkers (nodes), respectively. Both models require the following minimal ingredients: (1) preferential disassembly of edges or nodes under small tension or force, (2) a small but nonzero rate of random disassembly, and (3) energy injection upon assembly. Our models are robust to variations in random disassembly rates, can recover from drastic structural disruption, and exhibit diffusion of nodes and edges in steady state, displaying representational drift similar to that found in neuronal activities and physical learning circuits. We propose that the cortex is an example of “tunable matter,” *i.e*., its mechanosensitive remodeling dynamics tune properties of its edges and nodes so that the cortex as a whole can maintain robust but flexible rigidity in fluctuating mechanical environments.

## I. INTRODUCTION

The actin cortex is a dynamic network of actin filaments and actin-binding proteins that lies just beneath the plasma membrane. It plays a central role in shaping cells and regulating essential mechanical functions, including division, migration, and morphogenesis [1–4]. Having evolved under biological pressures that often require both robustness and adaptability, the cortex behaves very differently from engineered polymer networks [5]. It maintains *rigidity homeostasis*, i.e., a steady state of positive bulk and shear moduli, even as its architecture is constantly remodeled [5, 6]. During remodeling, filaments are pruned (removed by disassembly) and new ones are assembled, and crosslinkers are detached and reattached elsewhere. The fraction of unchanged filaments and crosslinkers decay exponentially with a time scale of tens of seconds [7–10]. The cortical structures undergo diffusion or superdiffusion [11–15], suggesting that remodeling processes leave no permanently persistent structural correlations. Despite this rapid turnover, the cortex exhibits striking long-lived mechanical memory: Following a step strain, stress decays algebraically as *σ*(*t*) ∼*t*^−*α*^ rather than exponentially [16–20], suggesting that memory of the deformed state survives long after the filaments and crosslinkers at the initial loading have been replaced. The cortex’s seemingly paradoxical behaviors of rigidity, remodeling, and elastic memory challenge conventional mechanical intuition, and raise the fundamental question of why the cell invests heavily in complete turnover [21], when a static or partially remodeling network might be sufficient to maintain rigidity.

Adding to this puzzle is the network structure of the actin cytoskeleton. In three dimensions, hyperstatic elastic networks with a mean coordination number (i.e., number of filaments joined at crosslinkers) above the Maxwellian threshold 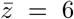 are inherently rigid [22]. While this is the case for cortices of certain embryonic [23] and red blood cells [24], the actin networks of many important cell types, including HeLa, have 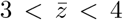 [25, 26], far below the Maxwellian threshold. Rigidity at such low coordinations can be attributed to the nonzero bending stiffness of actin filaments [27, 28]. However, tensegrity (tensional rigidity) can be an important contributor to overall rigidity [29, 30]; *i*.*e*., even in the absence of bending stiffness, prestressed filaments shift the rigid-to-floppy phase transition of an elastic network to 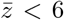 [31–33]. These prestresses may originate from internal contractile forces from myosin [4, 34] and are balanced by the attachment to the plasma membrane [35]. If cortex rigidity indispensably depends on filament tensions, it is even more surprising that rigidity persists robustly amid rapid remodeling, in which each filament pruning or crosslinker unbinding event leads to some tension loss. To understand rigidity homeostasis cleanly, we treat this most challenging case by neglecting bending rigidity.

In our previous work [36], we found that random pruning causes elastic networks to lose rigidity at biologically accessible prestrains (≲ 5%) even at 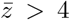. This result suggests that the cortex must effectively implement rules that selectively assemble or disassemble filaments and crosslinkers. Recent experiments indeed point to the presence of mechanosensitive remodeling: Actin filaments under lower tension are shown to be more susceptible to pruning by actin depolymerizing factor (ADF)/cofilin [37]. Common crosslinkers such as *α*-actinin [38] and vinculin [39] exhibit catch-bonding, i.e., they are more likely to unbind under smaller tensions.

Motivated by the above and recent agent-based computational studies [40–44], here we propose a resolution to the cortex’s paradoxical properties by viewing it as a mechanosensitive “tunable matter”. We model the actin cortex as a low-coordination elastic network consisting of spring edges (filaments) under tension joined by point-like nodes (crosslinkers and branchers). The edges and nodes are “tunable,” or individually adjustable to be there or not there according to local tensions, to achieve collective rigidity, much as parameters in neural nets are tuned so that the network as a whole can perform AI tasks. Unlike in conventional neural nets, where parameters are tuned using variants of gradient descent, edges/nodes in our models are tuned via *local mechanosensitive processes*. In that sense they are similar to physical Contrastive Local Learning Networks [45, 46] that use local rules to learn AI tasks.

As a first foray into the effects of mechanosensitive local rules on capturing the multitude of cortex-like behaviors, we have built two complementary minimal models. The first focuses on the dynamics of filaments with tension-inhibited pruning, and the second on the dynamics of crosslinkers with catch bonding. Our simulations reveal some general principles for networks under tensegrity to achieve rigidity homeostasis with complete turnover. Specifically, both models require all the following ingredients: (1) preferential disassembly of edges or nodes under small tension, (2) a small fraction of random disassembly, and (3) energy injection upon assembly that compensates for energy dissipation due to disassembly. Importantly, these ingredients appear to be minimal. The networks lose rigidity under purely random disassembly or if there is no energy injection, and fail to turn over under purely mechanosensitive disassembly. In other words, the three ingredients are necessary and sufficient for rigidity homeostasis with complete turnover.

Remarkably, elastic memory *emerges* from both minimal models: Like the cell cortex, our networks exhibit algebraic instead of exponential stress decay. Our simulations suggest that the origin for the elastic memory is that mechanosensitive disassembly preserves tension-weighted anisotropy. Many other emergent behaviors of our models also align with experimental observations, including diffusion of edges and nodes, heavy-tailed distribution of their displacements, the capability to recover from severely damaged states, and the closeness to a rigidity phase transition. These results support our hypothesis that the cortex is tunable matter that uses mechanosensitive local rules to explore the vast space of network architectures in order to achieve biologically desirable properties. While it is a longstanding view that turnover enables the actin cortex to respond effectively to its mechanical environment [47–49], our models formalizes this intuition through explicit biologically-plausible mechanisms, enabling quantitative exploration of cellular mechanical adaptability.

The rest of the paper is organized as follows. Section II describes our mechanosensitive models for filament and crosslinker dynamics. Section III demonstrates rigidity homeostasis with complete turnover. Section IV presents results for elastic memory. Sensitivity analysis of our models is given in Sec. V. Finally, Sec. VI provides concluding remarks.

## II. MODELS

Here we introduce our models with mechanosensitive edges and nodes, respectively, that generate remodeling elastic networks representing the actin cortex. Tension-rigidified networks are in the stretching-dominated “affine mechanical” regime, for which the ratio between bending and stretching contributions to the shear modulus is given by the volume fraction of actin filaments [50, 51]. The volume fraction of the actin cortex is reported to be 0.1–2% [52, 53], and thus can be assumed to be very small. We therefore neglect bending and focus on stretching.

For both models, we consider three-dimensional (3D) networks of harmonic springs joined by point-like nodes in a cubic box of volume *V* under periodic boundary conditions. The spring edges represent actin filaments and nodes joining the edges represent network-forming crosslinkers (e.g., filamin) or branching nucleators (e.g, Arp-2/3). We assume that the Young’s modulus of *Y* (with units of pressure) and the cross-sectional area *a* (with units of length squared) of the springs are constant and uniform across all springs, as they are set by the corresponding properties of actin filaments. Note that with this choice for units, the uniaxial strain on a filament is equivalent to its tensional force in units of *aY* . Furthermore, let *ℓ* be a characteristic mesh size (with units of length) such that *ℓ* ≫ *a*^1*/*2^. In what follows, the bulk and the shear moduli are measured in units of *Y*, the tension on filaments is measured in units of *aY*, and the elastic energy is measured in units of *aℓY* .

To reflect the fact that in the cortex, branchers and crosslinkers create degree-3 and -4 nodes, respectively, the degrees of all nodes are constrained so that *z*_*i*_ ≤ 4 throughout our simulated remodeling of the networks, where *z*_*i*_ is the degree of node *V*_*i*_. In each time step, changes to edges or nodes are performed as described in the subsections below, followed by a energy minimization via the Newton-Raphson method to equilibrate the physical degrees of freedom (the node positions) to satisfy force balance, before proceeding to the next iteration. That is, we work in the quasi-static limit by which it is assumed that force balance is much faster than structural turnover. For computational efficiency, we allow edges to pass through each other. However, we include Monte-Carlo based steric repulsion to compensate for the use of such “phantom” edges, as detailed in the subsections below.

For results in Sec. III, the initial configuration for both models is a network initially created from a jammed packing with 5% prestrain, pruned using the min-*E*_*i*_ rule described in Ref. [36] until all nodes are of degrees 4 or less. At that point the mean coordination number is 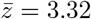. Other properties of this initial state, including the number of nodes and edges, are listed in Table I. Sensitivity analysis of our models to initial configurations is provided in Sec. V.

**TABLE I.** Parameters used in our network models for mechanosensitive edge and node dynamics for results in Sec. III.

| Property of jamming-derived initial state |  | Symbol | Value | Units |
| --- | --- | --- | --- | --- |
| Box side length | | $V^{1/3}$ | 9.477 | $\ell$ |
| Number of nodes | | $N_V$ | 1359 | - |
| Number of edges | | $N_E$ | 2030 | - |
| Mean node degree | | $\bar{z}$ | 3.32 | - |
| Rest length of all edges | | $l_{ij}$ | 0.85 | $\ell$ |

| Edge model parameter | Symbol | Value | Units | Node model parameter | Symbol | Value | Units |
| --- | --- | --- | --- | --- | --- | --- | --- |
| Threshold tension for pruning | $\Theta_E$ | 0.001 | $aY$ | Threshold tension for splitting | $\Theta_V$ | 0.001 | $aY$ |
| Random pruning events per iteration | $\kappa_E$ | 1 | - | Random splitting events per iteration | $\kappa_V$ | 0.5 | - |
| Nucleus concentration | $c_\nu$ | 0.5 | $\ell^{-3}$ | Capture radius | $r_c$ | 1.5 | $\ell$ |
| Insertion prestrain | $\epsilon$ | 0.05 | - | Insertion prestrain | $\epsilon$ | 0.12 | $aY$ |
| Maximum allowed rest length | $l_m$ | 3.0 | $\ell$ | | | | |
| Steric test points per nucleus | $n_s$ | $\geq 30$ | - | Steric test orientations per dangling edge | $n_s$ | $\geq 15$ | - |
| Steric sampling radius | $r_s$ | 1.0 | $\ell$ | Steric sampling radius | $r_s$ | 0.85 | $\ell$ |

### A. Mechanosensitive edge model

In our model for edge assembly and disassembly, the nodes are persistent and the edges are removed or inserted, representing pruning and assembly of filaments, respectively, to endow each edge with a binary tunable degree of freedom. As illustrated in Fig. 1(a), the key dynamic ingredients are tension-inhibited pruning, random pruning, and insertion of prestrained edges, through which elastic energy is injected into the network.

**FIG. 1.**
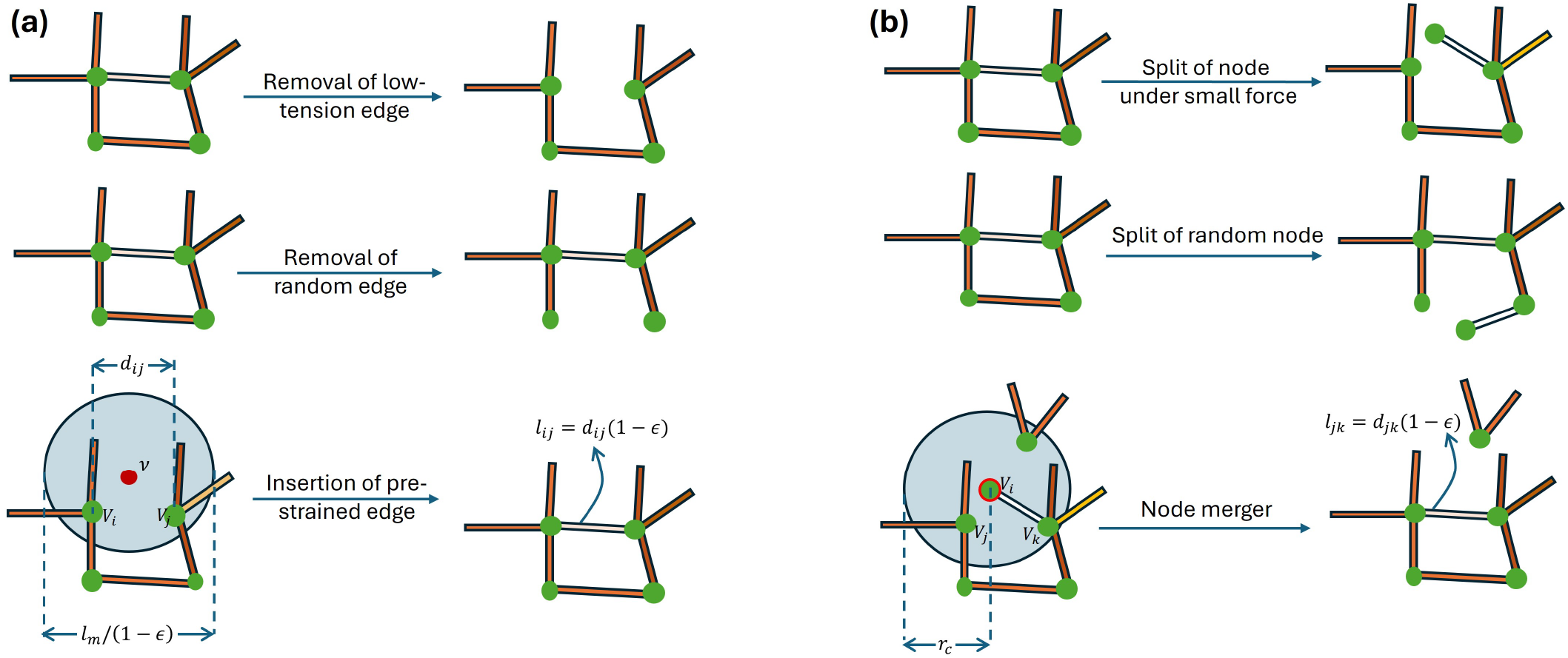
Illustrations of the dynamic ingredients our network models. (a) Elements of the mechanosensitive edge model. (Top) An edge under low tension is always removed. (Middle) An edge can be randomly removed with a small probability regardless of its tension. (Bottom) A prestrained edge is inserted between two nodes of degrees ≤ 3 in the neighborhood of a nuclei. (b) Elements of mechanosensitive node model. (Top) A node with an attached edge under small tension (therefore exerting a small force) always splits by detaching that edge. (Middle) A node of degree ≥ 2 can split randomly regardless of the tensions on its associated edges. (Bottom) A degree-1 node merges with its nearest neighboring node within a capture sphere. Upon a merger, the rest length of the dangling edge is adjusted to impose an edge prestrain *ϵ*. We implement steric-biased edge insertions and node mergers to achieve homogeneous steady states, but steric repulsion is not needed for rigidity homeostasis.

In each iteration, all edges with tensions smaller than a threshold Θ_*E*_ are removed [Fig. 1(a), Top]. In addition to this mechanosensitive (tension-inhibited) pruning, exactly *κ*_*E*_ = 1 edge per iteration is randomly chosen to be removed regardless of its tension [Fig. 1(a), Middle]. To insert edges into the network, we first generate *c*_*v*_ *V* points in the simulation box that represent nuclei [red point in Fig. 1(a, Bottom)], where *c*_*v*_ is the nucleus concentration. Due to steric repulsion, we assume that new filaments are preferentially formed at locations with lower local density of existing filaments. As such, the positions of the nuclei are chosen according to a Monte Carlo based steric rule: To generate each one of the nuclei, we first sample *n*_*s*_ “test points” from the uniform distribution in the box to probe local filament densities. The test point with the fewest edge midpoints in a steric sampling window of radius *r*_*s*_ = *ℓ* is chosen to be the new nucleus; *c*_*v*_ *V* of such nuclei are generated in this manner. We verified that the results in Sec. III A are insensitive to *n*_*s*_ as long as *n*_*s*_ ≥ 30. In the Appendix, Sec. C, we will show that the steric rule is not required to sustain rigidity, but is merely needed to maintain a spatially *homogeneous* steady state.

For each nucleus *v*, the candidate nodes to be connected by the newly formed edge are those with *z*_*i*_ < 4 that lie within the sphere of radius *l*_*m*_*/*[2(1 − *ϵ*)] centered at *ν*[shown by the blue region in Fig. 1(a), Bottom], where *l*_*m*_ is the maximum allowed rest length and *ϵ* is the insertion prestrain. Among these candidate nodes, two nodes (denoted *V*_*i*_ and *V*_*j*_) are randomly selected, and a new edge is inserted between them with rest length *l*_*ij*_ = *d*_*ij*_(1 − *ϵ*), where *r*_*ij*_ is the Euclidean distance between *V*_*i*_ and *V*_*j*_ [Fig. 1(a), Bottom]. If there are fewer than two nodes with *z*_*i*_ < 4 in the aforementioned nucleation sphere, no new edge is inserted.

During the simulation, isolated (degree-0) nodes are occasionally generated, i.e., when all the originally attached edges have been removed. In such cases, the isolated node is displaced to the nearest edge midpoint, and the destination edge “breaks” into two edges joined by the incoming node, which now has *z*_*i*_ = 2. That is, we assume that standalone crosslinkers and branchers rapidly bind to nearby existing filaments. They thus join the population of *z*_*i*_ < 4 nodes that could be connected by new edges. This “hopping” treatment facilitates the addition of filament-node links but makes the network slightly weaker because an initially straight edge becomes two edges joined by a “hinge” node without a bending penalty. In our simulations (Sec. III A), we find that on average there are only 0.22 “hopping” isolated nodes per iteration, out of the total 1359 nodes. Thus, hopping isolated nodes are insignificant to the evolution trajectory of the network.

### B. Mechanosensitive node model

In our model for node dynamics, the edges are persistent, while the connectivities *z*_*i*_ of the nodes evolve via splits and mergers of nodes, and their rest lengths change according to the prestrain parameter *ϵ*. As illustrated in Fig. 1(b), the key dynamic ingredients are splitting of nodes under small pulling forces, random node splitting, and node merging upon which energy is injected. We note that while the binding and unbinding of a crosslinker or brancher can correspond to either *z*_*i*_ → *z*_*i*_ ± 2 or *z*_*i*_ → *z*_*i*_ ± 1, we consider here only the latter case for simplicity.

In each iteration, consider a random edge *E*_*α*_ connected to node *V*_*i*_ with *z*_*i*_ ≥ 2. To implement the catch-bonding behavior, if the tension on *E*_*α*_ is smaller than a threshold value Θ_*V*_ (thereby exerting a small force on *V*_*i*_), then *V*_*i*_ is split by detaching *E*_*α*_, forming two new nodes with degrees *z*_*i*_ → *z*_*i*_ −1 and 1, respectively. To implement random node splitting, in each iteration, with probability *κ*_*V*_ = 0.5, a node with *z*_*i*_ ≥ 2 is randomly chosen to split by detaching one of its edges, also chosen randomly. A dangling edge generated by node splits is allowed to freely rotate about its other node connected to the rest of the network. However, to compensate for the use of “phantom” edges and to create larger angles between edges on a node, an orientation with smaller steric repulsion is selected, again according to a Monte Carlo based steric rule: For each dangling edge, *n*_*s*_ orientations are randomly sampled, and the orientation that gives the fewest edge midpoints in a sampling sphere of radius *r*_*s*_ ≈ *ℓ* centered at the dangling (degree-1) node is chosen to be the adopted orientation prior to node mergers. Simulation results in Sec. III B are insensitive to *n*_*s*_ as long as *n*_*s*_ ≥ 15.

To mimic crosslinker binding, each degree-1 node *V*_*i*_ merges with its nearest node *V*_*j*_ with *z*_*j*_ ≤ 3, provided that *V*_*j*_ lies within the “capture sphere” centered at *V*_*i*_ with radius *r*_*c*_, i.e., *d*_*ij*_ ≤ *r*_*c*_. If there is no candidate node with *z*_*j*_ ≤ 3 in the capture sphere, no merger is performed. To inject elastic energy into the network upon assembly, the rest length of the corresponding dangling edge is modified to *l*_*ik*_ = *d*_*jk*_(1 −*ϵ*) prior to relaxation, where *V*_*k*_ is the neighboring node of *V*_*i*_; see Fig. 1(b), bottom panel. Table I summarizes the parameters of our models, as well as their values used to produce the results for rigidity homeostasis shown in Sec. III.

It is important to remark on the assumption about time scales built into our models. Firstly, by implementing the quasistatic algorithm, in which we relax the networks to the energy equilibrium at every time step, we have assumed that force equilibration is much faster than processes that modify the network architecture, i.e., filament pruning and creation, and crosslinker binding and unbinding. This assumption is valid, as the cytoskeleton without architectural change behaves like a poroelastic medium [54]. The poroelastic relaxation timescale is given by *τ*_*p*_ = *ηL*^2^*/*(*Eξ*^2^), where *η* ≈ 10^−3^ Pa·s is the cytosol viscosity, *E* = 10^3^ Pa is the Young’s modulus of the actin network, *ξ* = 50 nm is the characteristic mesh size and *L* = 1*µ*m is the characteristic length scale of deformation. According to this estimate, *τ*_*p*_ is 0.4 millisecond, much smaller than the cytoskeleton’s remodeling length scale, which is on the order of tens of seconds. This separation of time scales is crucial for mechanosensitivity and has been reported in several experiments [55, 56]. Secondly, we prestrain the filaments immediately upon assembly as the simplest way to inject energy, which does not require explicit incorporation of myosin motors. With this treatment, the energy injection time scale is assumed to be much faster than turnover time scale, a valid assumption as the timescale for actomyosin power strokes is on the order of a millisecond [57]. The only time scale hard coded into our models is now the inverse of the random disassembly rates, 1*/κ*_*E*_ and 1*/κ*_*V*_ . While it is in general difficult to relate these time scales of our pseudodynamics to real time, once a steady state is reached, the emergent turnover time scale can be related to the experimentally observed values, i.e., tens of seconds.

For model validation, we confirmed that our algorithm for linear response (described in Appendix A) generates known results for the diamond and the cubic elastic networks in Ref. [58], and that our random and tension-inhibited disassembly algorithms reproduce the elastic moduli in our earlier paper [36]. Note that in both models, nodes of degrees 0, 1, 2 serve only a bookkeeping role as part of the candidate pools for nodes that can have edges added to them or that can merge with other nodes. When reporting the mean coordination number 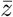, only nodes of degrees 3 and 4 (physical ones) are counted.

## III. RESULTS FOR RIGIDITY HOMEOSTASIS

In this section, we present results for rigidity homeostasis for both models. We show that our models capture the minimal requirements for rigidity homeostasis, in the sense that omission of any of dynamical ingredients illustrated in Fig. 1 (*i*.*e*., tension- or force-sensitive disassembly, random disassembly, and energy injection upon assembly) causes the networks either to lose rigidity or fail to turn over completely.

### A. Results for the mechanosensitive edge model

Figure 2 shows results for rigidity homeostasis in the mechanosensitive edge model (Sec. II A), using the parameter values given in Table I. The network reaches a dynamic steady state, for which a representative snapshot is shown in Fig. 2(a) and a video showing the evolution of the network configuration at the steady state is given in the Supplemental Information (SI, Ref. 59), Video S1. At the steady state, the mean node degree is 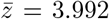, which is larger than that in the initial state (Table I), indicating that almost all nodes have reached the saturation coordination number of 4 due to the high nucleus concentration. The colors of edges in Fig. 2(a) represent their strains, (or equivalently, tensions in units of *aY* ). The edge tension distribution of the configuration in Fig. 2(a) is given in Fig. 9 (solid curve) in the Appendix B. The mean strain is 0.85%, and the maximum strain is 4.8%, just below the insertion prestrain *ϵ* = 5%. Note that the mean tension is much smaller than *ϵ*. Indeed, we find that despite our 5% insertion prestrain, the tensions on many newly inserted edges decrease significantly after energy minimization, and are quickly recycled by tension-inhibited pruning. For example, in the first 200 iterations, there are in total 1881 edge insertions, but only 780 of these new edges survive at *t* = 200.

**FIG. 2.**
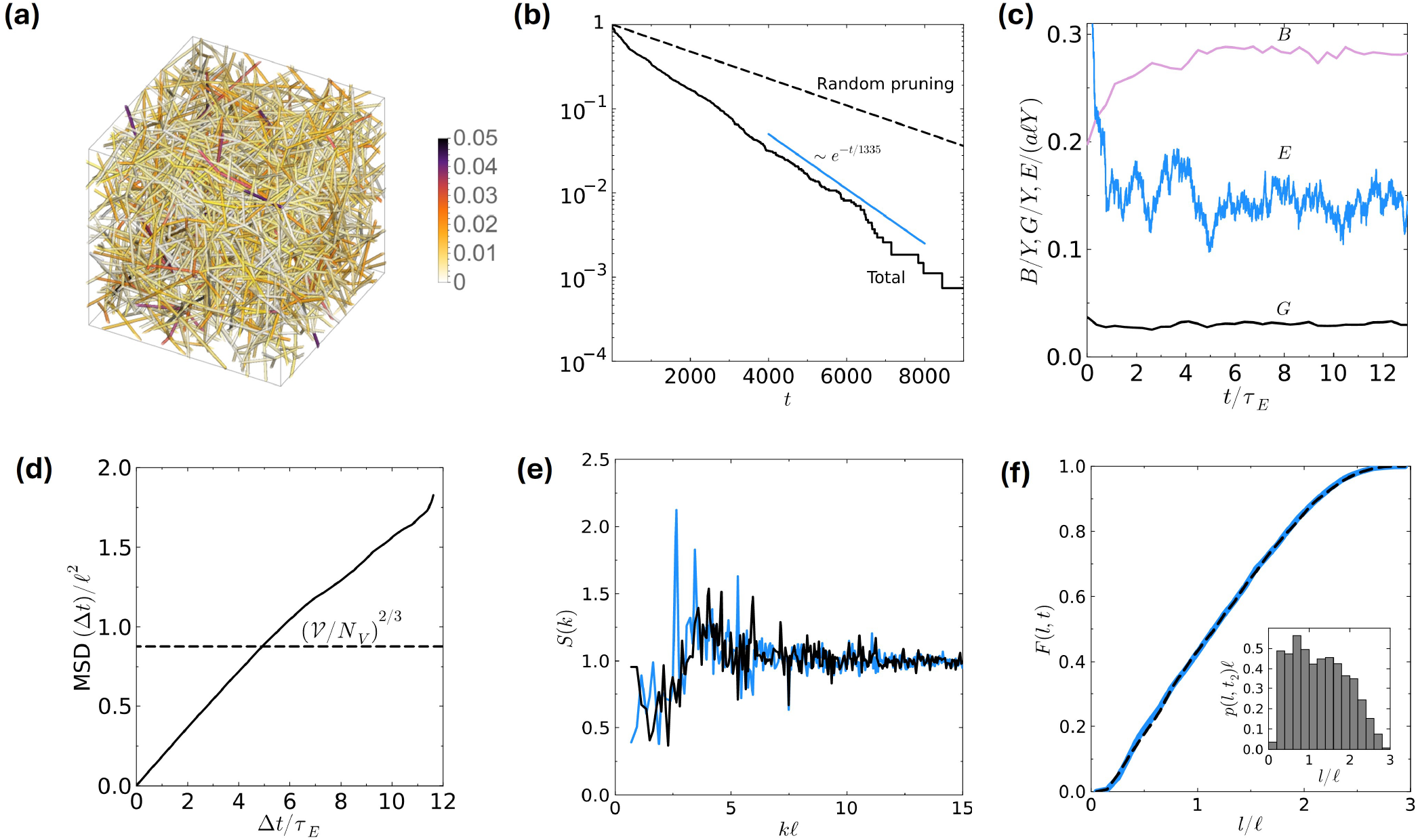
Rigidity homeostasis in mechanosensitive edge model described in Sec. II A. (a) Sample snapshot of the network in its dynamic steady state (*t* = 6300). The color bar indicates edge tensions. (b) Measurement of edge turnover. The solid curve shows the fraction of edges *γ*_*E*_ (*t*) in the initial configuration that remain unpruned at each iteration *t*. The dashed curve shows edge turnover contributed by random pruning. (c) Evolution of the bulk modulus *B*, the shear modulus *G*, and the elastic energy *E*. These quantities clearly reach steady states. (d) Time-averaged mean squared displacement of nodes. Nodes that have attained zero-degree status during the interval between *t* and *t* + Δ*t* are not included in the MSD calculations. (e) Structure factors for edge midpoints averaged over configurations in iteration intervals *T*_1_ = [7000, 9000] (blue) and *T*_2_ = [14000, 16000] = *T*_1_ + 5.2*τ*_*E*_ (black). (f) Cumulative edge length distribution functions *F* (*l, t*) at iterations *t*_1_ = 7000 (blue) and *t*_2_ = 14000 = *t*_1_ + 5.2*τ*_*E*_ (black, dashed). The inset shows the edge length probability distribution function *p*(*l, t*) at *t*_2_ = 14000.

To study the turnover of edges quantitatively, we plot in Fig. 2(b) the fraction of edges in the initial state that has *not* been pruned at iteration *t*, denoted *γ*_*E*_(*t*), shown by the solid curve. It is clear that this fraction decays exponentially with *t* at large *t*, i.e., *γ*_*E*_(*t*) ∝ exp(−*t/τ*_*E*_), with a characteristic turnover time scale *τ*_*E*_ = 1335 iterations. The dashed curve represents the hypothetical edge turnover if there were merely *κ*_*E*_ random pruning events and *κ*_*E*_ insertions in each iteration, given by

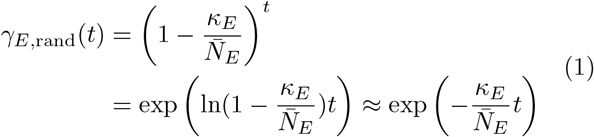

This curve has a characteristic turnover time scale 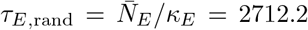, and is an upper bound to *γ*_*E*_(*t*) because it does not include tension-inhibited pruning. By comparing the turnover times for the two cases, we see that tension-inhibited pruning approximately doubles the rate of edge turnover compared to merely random pruning. Yet, we find that in each iteration, there are on average 8 times more tension-inhibited pruning events than random pruning events. The fact that the net turnover rates only increases by a factor of 2 (rather than 8) again indicates that tension-inhibited pruning primarily acts on newly inserted edges that have relaxed their tensions upon energy minimization.

Fig. 2(c) shows the evolution of linear elastic properties of the network, including the bulk modulus *B* and shear modulus *G*, as well as the total elastic energy *E*. These quantities reach steady state by ∼ *t* = 7000 ≈ 5.2*τ*_*E*_. Statistical tests confirming that a steady state is reached are described in SI, Sec. 1, where we show that for two time windows separated by 5.2*τ*_*E*_, *B, G, E*, 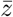 and the edge tension distributions are statistically indistinguishable according to two-sample t-tests and the Kullback-Leibler divergence [60]. Interestingly, in the transient phase before the steady state is reached, the elastic energy decreases while the bulk modulus increases. The shear modulus stays approximately constant throughout. This increase in *B* despite decreasing *E* is due to the fact that all inserted edges have identical insertion prestrain *ϵ* = 0.05, whereas edges under very low and very high tensions are gradually pruned via tension-inhibited and random pruning, respectively. These processes result in a more uniform edge tension distribution as the network evolves (see Appendix B). Uniformity in tension distribution is known to increase *B* at comparable mean edge tensions [61, 62].

To elucidate the evolution in microstructure and the displacement of network elements, we plot in Fig. 2(d) the time-averaged mean-squared displacements (MSD) of nodes. Note that when computing MSD(Δ*t*), we have excluded nodes that have become degree-0, because we aim to determine node displacements due to pruning and insertion dynamics of edges alone, whereas “hopping” of degree-0 nodes is manually imposed and does not arise from such dynamics. Figure 2(d) shows that the MSD increases linearly with Δ*t* and grows beyond the square of the characteristic separation between nodes (*V /N*_*V*_ )^2*/*3^ = 0.878*ℓ*^2^. According to the Lindemann criterion [63], a solid should lose its rigidity once the length scale for fluctuations exceeds some fraction of this distance. Furthermore, we compute the node displacement fluctuation characterized by the self van Hove function, which is the time-averaged probability distribution of particle displacements that take place during the time interval (*t, t* + Δ*t*). We find that the self van Hove function does not obey a Gaussian distribution as one might expect for diffusive behavior. The distribution is heavy-tailed [Fig. 3(a)], consistent with experimental findings [15]. It is well fitted by a truncated Lorentzian; see Appendix C.

**FIG. 3.**
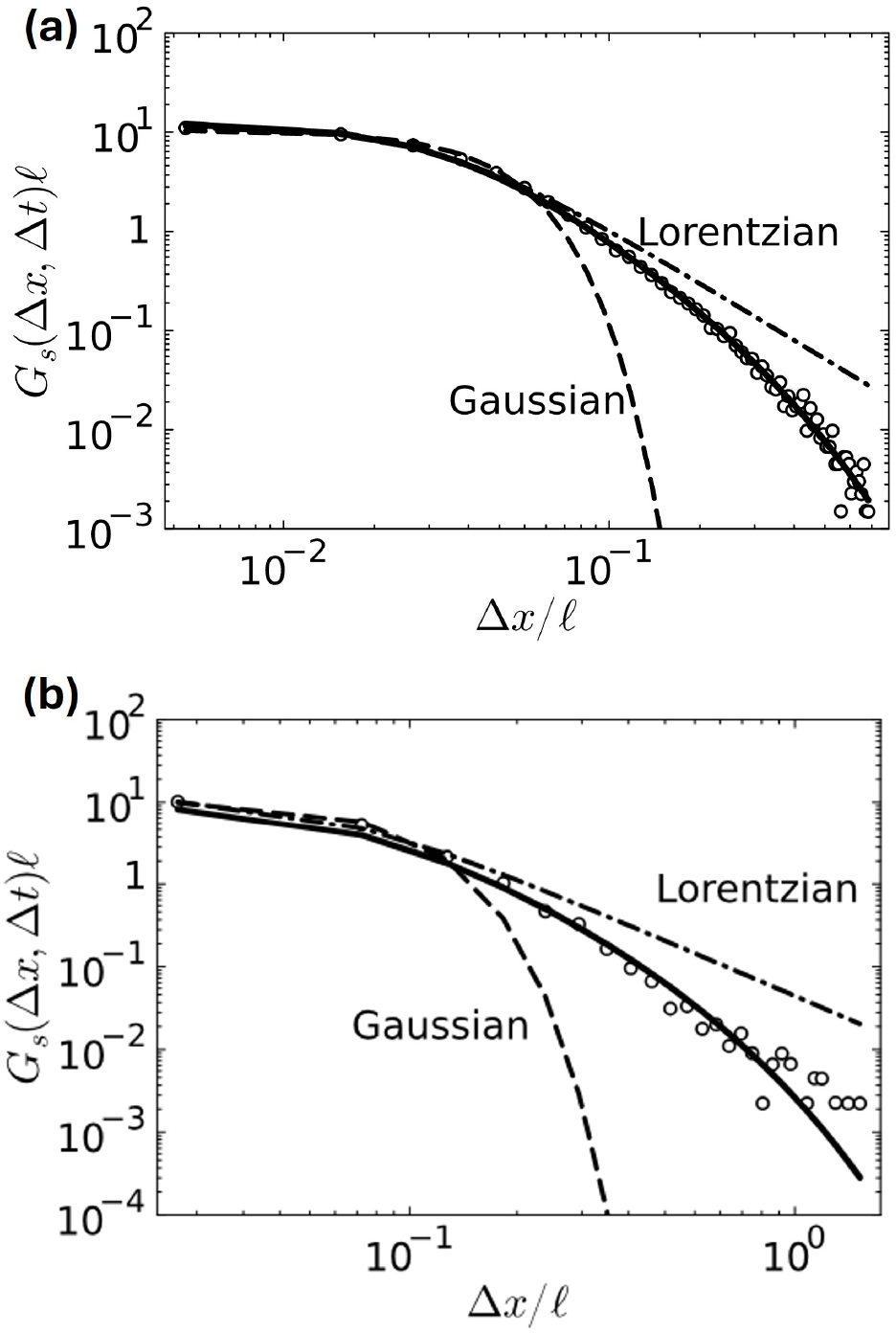
Simulated self van Hove functions *G*_*s*_(Δ*x*, Δ*t*) of (a) nodes in the edge model with time interval Δ*t* = 50 and edges midpoints in the node model with Δ*t* = 50 (circles). Both functions are heavy-tailed and can be accurately described by exponential-truncated Lorentzian distributions, Eq. (14), shown by the solid curves. The best-fit Gaussian (dashed) and non-truncated Lorentzian (dash-dotted) are also shown.

To investigate whether statistical properties of the network architecture are invariant at the steady state, we compute the time-averaged structure factors *S*(*k*) of edge midpoints in two time windows *T*_1_, *T*_2_ separated by 5.2*τ*_*E*_, *i*.*e*., *T*_1_ = [7000, 9000] and *T*_2_ = [14000, 16000], with a configuration snapshot taken every 50 iterations [Fig. 2(e)]. The structure factor for a point configuration is a commonly used microstructural descriptor to characterize the magnitude of density fluctuation at wavevector *k* and is the Fourier representation of the pairwise correlations of points in reference to a Poisson point process [64]. We also compute the cumulative distribution functions 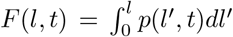 of edge rest lengths at *t*_1_ = 7000 and *t*_2_ = 14000 [Fig. 2(f)], where *p*(*l*^*′*^, *t*) is the associated edge-length probability distribution function, whose form at *t*_2_ is shown in the inset. Clearly, the structure factor and the edge-length distribution are statistically invariant. Taken together, the results in Fig. 2 show that our model achieves rigidity homeostasis with complete edge turnover while maintaining a dynamic steady state in terms of elastic moduli and microstructural statistics.

To show that all ingredients depicted in Fig. 1(a) are necessary for rigidity homeostasis, we have also studied two control models, one without random pruning (i.e., *κ*_*E*_ = 0) and the other without tension-inhibited pruning (i.e., Θ_*E*_ = 0). All other details and parameters are identical to those described in Sec. II A and Table I. Our results are summarized in Fig. 4, where we plot the fraction of unchanged edges, *γ*_*E*_(*t*) on the left axis and the elastic energy *E* on the right axis. The curve for *γ*_*E*_(*t*) shows that without random pruning, one cannot achieve complete turnover. Indeed, *γ*_*E*_(*t*) stalls at 93% at *t* = 54, at which point the network stops evolving because all edges have tension greater than Θ_*E*_ and no new edge can be inserted, because any two nodes with *z*_*i*_ ≤ 3 are separated by a distance greater than *l*_*m*_*/*(1− *ϵ*). On the other hand, the dashed curve for *E* shows that without tension-inhibited pruning, the elastic energy decays to zero, *i*.*e*., the network loses rigidity. Comparing the curves for *E* in Figs. 2(c) and 4, it may seem counterintuitive that removing more edges per iteration through tension-inhibited pruning *preserves rigidity*. This phenomenon is a result of our constraint that *z*_*i*_ ≤ 4 on all nodes. Tension-inhibited pruning creates more candidate nodes with *z*_*i*_ ≤ 3 that can be joined by edges with sufficiently high tension to remain, while random pruning leads to avalanches that rapidly degrade rigidity [36].

**FIG. 4.**
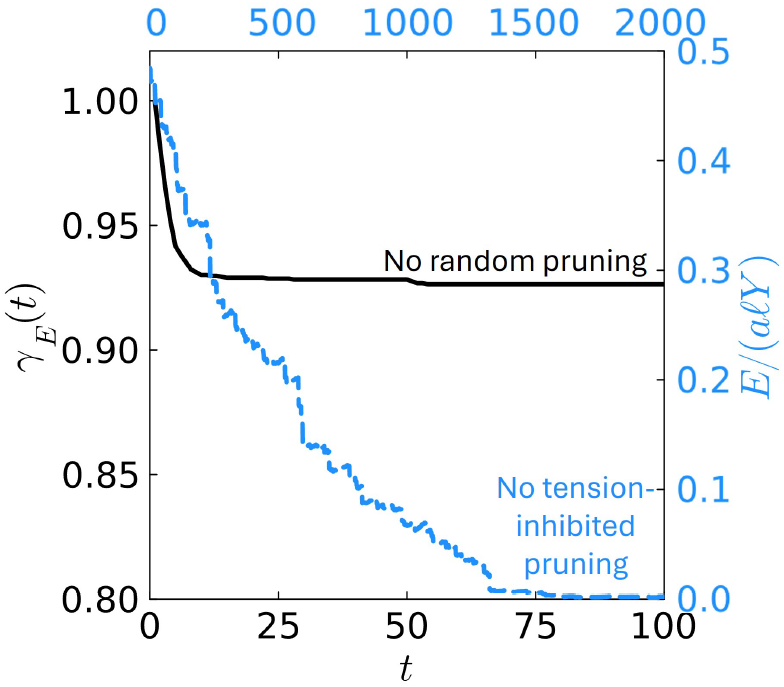
(Black, solid) Turnover of edges without random pruning, (i.e., *κ*_*E*_ = 0). The network stops evolving when there are still *γ*_*E*_ = 92.6% unpruned edges. (Blue, dashed) Evolution of elastic energy *E* without tension-inhibited pruning, (i.e., Θ_*E*_ = 0). The network gradually loses rigidity.

### B. Results for the mechanosensitive node model

Figure 5 presents results for our mechanosensitive node model, described in Sec. II B, starting from the jamming-derived initial state and using the parameter values listed in Table I. Again, statistical tests confirm that a steady state is reached (SI, Sec 1). The steady-state characteristics 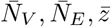, *B, E, G* are listed in Table II, Case 2. A representative snapshot of the network at the steady state is shown in Fig. 5(a) and a video is given in the SI, Video S2 [59].

**TABLE II.** Steady-state topological and mechanical properties of our models with different initial states and random disassembly rates. Simulation parameters except for *κ*_*E*_ and *κ*_*V*_ are kept identical to those in Table I.

| Case | Model | Rand. disassembly rate | Initial state | $\bar{N}_V$ | $\bar{N}_E$ | $\bar{z}$ | $B/Y$ | $G/Y$ | $E/(a\ell Y)$ |
| --- | --- | --- | --- | --- | --- | --- | --- | --- | --- |
| 1 | Edge | $\kappa_E = 1$ | Jamming | 1359 | 2712.2 | 3.992 | $0.286 \pm 0.002$ | $0.031 \pm 0.001$ | $0.14 \pm 0.01$ |
| 2 | Node | $\kappa_V = 0.5$ | Jamming | 1054.5 | 2030 | 3.950 | $0.35 \pm 0.01$ | $0.046 \pm 0.001$ | $0.37 \pm 0.04$ |
| 3 | Edge | $\kappa_E = 1$ | Fig. 5(a) | 1048 | 2087.9 | 3.988 | $0.224 \pm 0.006$ | $0.024 \pm 0.002$ | $0.10 \pm 0.01$ |
| 4 | Node | $\kappa_V = 0.5$ | Fig. 2(a) | 1380.6 | 2714 | 3.974 | $0.42 \pm 0.02$ | $0.067 \pm 0.003$ | $0.62 \pm 0.02$ |
| 5 | Edge | $\kappa_E = 1.5$ | Jamming | 1359 | 2708.4 | 3.989 | $0.292 \pm 0.005$ | $0.029 \pm 0.001$ | $0.12 \pm 0.01$ |
| 6 | Node | $\kappa_V = 0.75$ | Jamming | 1073.2 | 2030 | 3.935 | $0.345 \pm 0.008$ | $0.045 \pm 0.002$ | $0.36 \pm 0.04$ |
| 7 | Edge | $\kappa_E = 1$ | Diamond | 1359 | 2712.3 | 3.993 | $0.282 \pm 0.002$ | $0.033 \pm 0.001$ | $0.16 \pm 0.02$ |

**FIG. 5.**
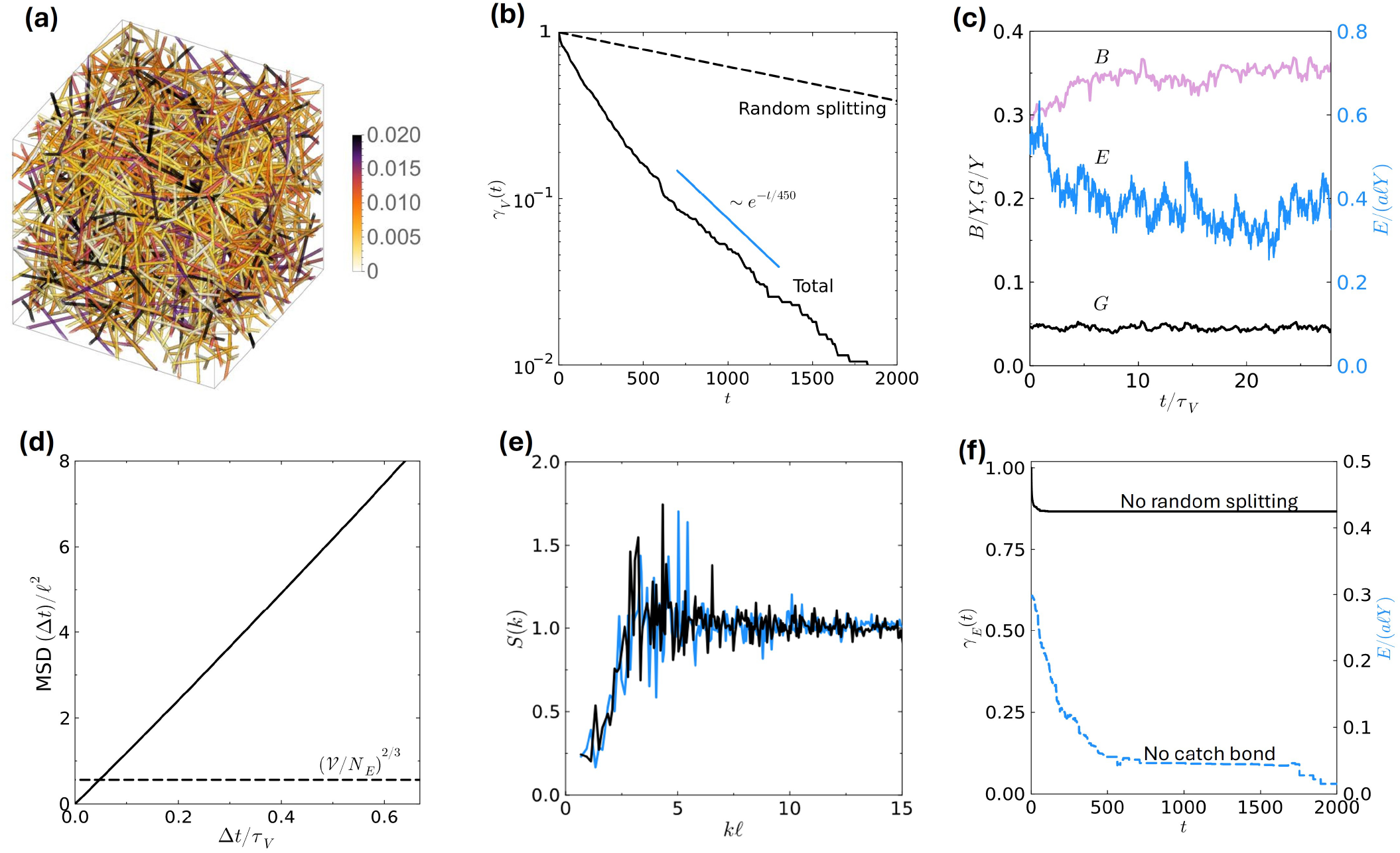
Rigidity homeostasis in mechanosensitive node model described in Sec. II B. (a) Sample snapshot of the network in its dynamic steady state (*t* = 10000). The color bar indicates the edge tensions. (b) Measurement of node turnover. The solid curve shows the fraction of nodes *γ*_*V*_ (*t*) in the initial configuration that have not changed coordination number at each iteration *t*. The dashed curve shows node turnover contributed by random splitting. (c) Evolution of the bulk modulus *B*, the shear modulus *G*, and the elastic energy *E*. These quantities clearly reach steady states. (d) Time-averaged mean square displacement of edge midpoints. (e) Structure factors for edge midpoints averaged over configurations in iteration intervals *T*_1_ = [7000, 7500] (blue) and *T*_2_ = [12000, 12500] = *T*_1_ + 11*τ*_*V*_ (black). (f) (Black, solid) Turnover of nodes in the model described in II B but without random splitting, (i.e., *κ*_*V*_ = 0). The network stops evolving when there are still *γ*_*V*_ = 86.6% unchanged nodes. (blue, dashed) Evolution of elastic energy *E* without catch bonds, (i.e., Θ_*V*_ = 0). The network gradually loses rigidity.

Results in Fig. 5 shows that the qualitative behaviors of the node model closely parallel those of the edge model. As shown by the solid curve in Fig. 5(b), the fraction of nodes that have never changed their degree up to iteration *t*, denoted *γ*_*V*_ (*t*), decays exponentially at large *t*, with a characteristic turnover time scale *τ*_*V*_ = 450 iterations. The dashed curve in Fig. 5(b) shows node turnover contributed by random splitting,

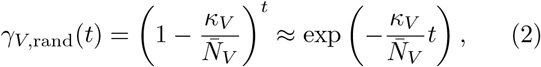

with a characteristic turnover time scale 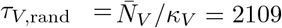. Thus, the inclusion of catch-bond splitting causes the turnover rate to roughly quadruple compared to if there were merely random splitting. Figure 5(c) shows that the bulk and shear moduli as well as the elastic energy reach steady states, confirming that this model achieves rigidity homeostasis. Due to the rotation of dangling edges and the change of their rest lengths, the edges undergo rapid diffusion, and the MSD of edge midpoints increases beyond the squared characteristic separation between edge midpoints (*V /N*_*E*_)^2*/*3^ = 0.560*ℓ*^2^ in less than 0.1*τ*_*V*_, as shown in Fig. 5(d). Again, we observe a heavy-tailed self van Hove function of edge midpoints that is well fitted by a truncated Lorentzian [Fig. 3(b)]. Finally, similar to the edge model, the structure factor of edge midpoints is statistically invariant. As shown in Fig. 5(e), the averaged *S*(*k*) in two time windows *T*_1_ = [7000, 7500] and *T*_2_ = [12000, 12500] = *T*_1_ + 11*τ*_*V*_ agree well with one another.

Despite the similarities between the results of our edge and node models, it is important to remark that edge turnover [Fig. 2(b)] and node diffusion [Fig. 2(d)] in the former model proceed much slower than node turnover [Fig. 5(b)] and edge diffusion [Fig. 5(d)], respectively, in the latter model. This is due to the fact that dangling edges are allowed to freely rotate in the node model, causing rapid increase in the MSD of edge midpoints. On the other hand, in the edge model, node displacements are solely caused by the fluctuating forces exerted by the filaments, and thus the evolution of the network microstructure is thus much slower. We have also computed MSD for edges that have not undergone rest length changes in the node model. We find that these edges also undergo diffusion, but at a much lower rate: Their MSD crosses the mesh size at *t* ≈ *τ* .

Again, we have verified that all dynamic ingredients shown in Fig. 1(b) are necessary for our node model to attain rigidity homeostasis. Indeed, Fig. 5(f) shows that without random splitting, (i.e., *κ*_*V*_ = 0), node turnover *γ*_*V*_ (*t*) stalls at 86.6%. On the other hand, without catch-bond splitting, (i.e., Θ_*V*_ = 0), the network loses rigidity in 2000 iterations.

## IV.RHEOLOGY AND ELASTIC MEMORY

An overcoordinated network can maintain rigidity under turnover, without any mechanosensitivity. Thus, here we consider the question: What advantage might be conferred by the strategy studied here, of a prestrained undercoordinated network 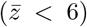 subject to mechanosensitive tuning?

Here, we show that one potential advantage is longlasting elastic memory. Cells are known to exhibit slow stress decay upon an imposed step strain that is consistent with a power law over the range measured. We hypothesize that mechanosensitivity allows the cortex to attain elastic memory well beyond the turnover timescale.

To investigate the response of our models upon an imposed step strain, we apply a pure shear to the steadystate snapshots of both models by deforming the simulation box basis with the strain tensor

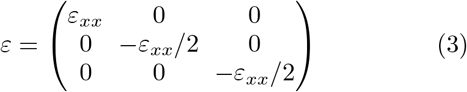

Following this deformation, we compute the stress relaxation modulus *G*(*t*) by monitoring the anisotropy of axial stresses: *G*(*t*) = [*σ*_*xx*_(*t*) − (*σ*_*yy*_(*t*) + *σ*_*zz*_(*t*))*/*2]*/*(3*ε*_*xx*_), where *t* is the timestep after the application of the strain. Such “simulated rheology” generate noisy stress curves, and thus we averaged over 60 and 120 trajectories for the edge and node models, respectively.

The simulated rheology reveals striking evidence of elastic memory. As shown by the scattered points in Fig. 6, *G*(*t*) are characterized by a transient fast decay followed by a slowly decaying tail that retains 10–20% of *G*(*t* = 0) even after the original crosslinkers have undergone 10*τ*, at which *γ*(10*τ* ) = 4.5 × 10^−5^ of the edges or nodes at *t* = 0 remain unchanged.

**FIG. 6.**
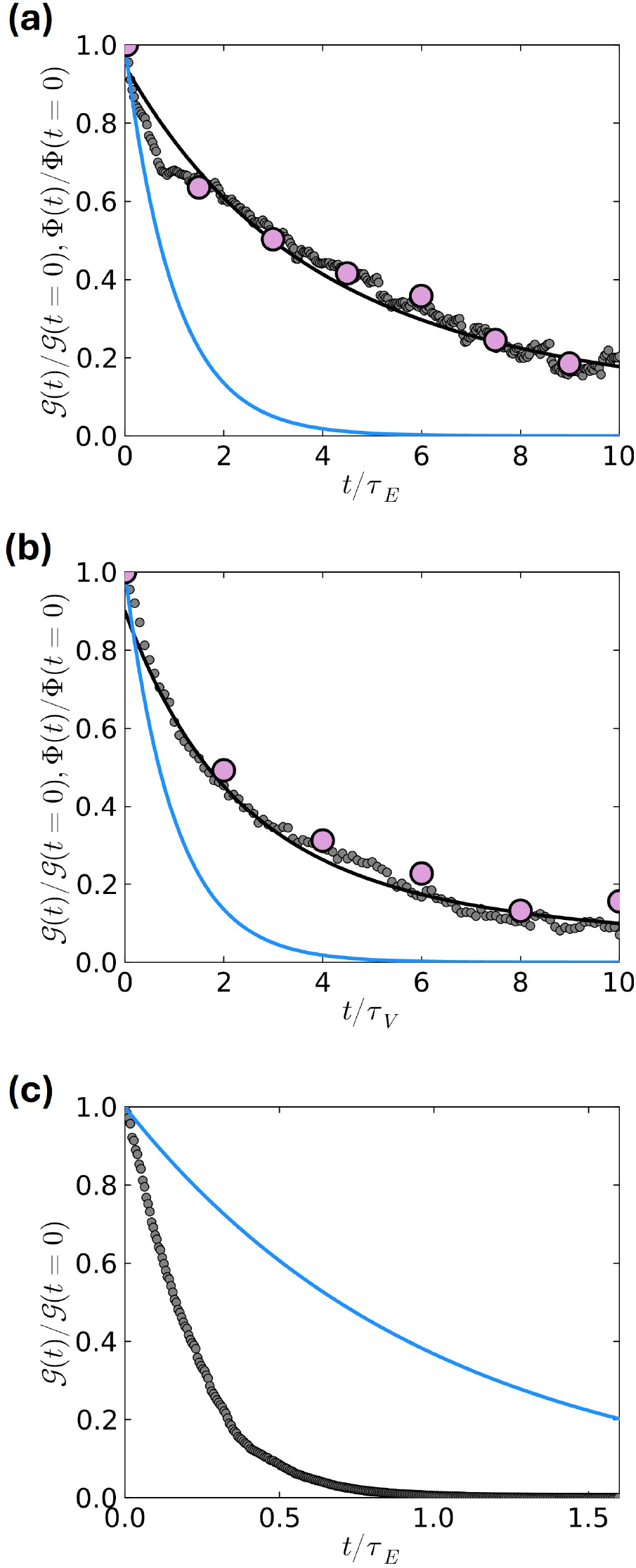
Normalized stress relaxation modulus *G*(*t*)*/ G*(*t* = 0) of the node model following a 2% pure shear step strain for (a) Edge model, (b) node model, and (c) overcoordinated control model. The gray points are from stress decay upon a pure shear deformation of simulation box. The black curves are *G*(*t*) computed from microrheology. The blue curves represent the corresponding filament and crosslinker turnover. The triangles represent the normalized anistropy order parameter (4) Φ(*t*)*/*Φ(*t* = 0)

To find the long-time behavior of (*t*), we use microrheology techniques, which can generate highly precise rheology data from the microscopic fluctuations of edge and node positions and the corresponding stress fluctuations in the undeformed simulation box; see details in Appendix B. The analytical (*t*) obtained from microrheology (Fig. 6, black curve) agree very well with our simulated rheology, and their fitted functional forms have a algebraic tail *t*^−1^ (Appendix B), further confirming that our model has an elastic memory well beyond *τ*_*E*_ and *τ*_*V*_ .

To demonstrate the role of mechanosensitivity in elastic memory, we investigate the mechanical response of an overcoordinated control model that maintains rigidity homeostasis with complete turnover, but does not have mechanosensitivity. Starting from a cubic grid network with *N*_*V*_ = 1000 (*z* = 6 for all nodes), we draw random pairs of nodes without replacement and join them by additional edges, until *z* = 7 for all nodes. To implement remodeling, we use degree-preserving edge swaps: two edges joining (*i, j*) and (*k, l*) are drawn randomly and replaced by edges joining (*i, l*) and (*k, j*). The rest lengths of the new edges are equal to the Euclidean distance between the nodes to be joined. Energy minimization is performed after every edge swap. Note that an overcoordinated network is used because an undercoordinated one would quickly lose rigidity with this random remodeling process. We apply the pure shear strain (3) to this model and monitor the stress relaxation and turnover of edges. The stress relaxation is found to be even faster than edge turnover, as shown in Fig. 6(c). Therefore, it is evident that mechanosensitive remodeling is crucial in realizing elastic memory.

To investigate the structural origin of the elastic memory, we compute an order parameter that measures tension-weighted anisotropy along the basis axes:

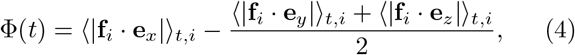

where **f**_*i*_ is the force on an edge *i*, **e**_*x*_ is the unit vector along the *x*-axis, and the averages ⟨⟩_*t,i*_ are taken over all edges and all trajectories at time *t* after the deformation. The values of the normalized anisotropy parameter Φ(*t*)*/*Φ(*t* = 0) are shown as triangle symbols in Fig. 6, and they closely match the trend of the stress decay. The slow decay of Φ(*t*) is expected: Upon applying the pure shear strain (3), edges aligned along the stretched direction **e**_*x*_ are under higher tension than edges along **e**_*y*_ and **e**_*z*_, and consequently disassembled at a slower rate. Thus, we have shown that the elastic memory is achieved through the remodeling process that preserves tension-weighted anisotropy.

## V. SENSITIVITY ANALYSIS

The actin cortex is known to exhibit remarkable robustness against external and internal stresses, local perturbations in network structure, and fluctuations in filament, crosslinker, and severing protein concentrations. Here, we examine the sensitivity of our models to three key factors: the initial state, the random disassembly rate, and the tension threshold for mechanosensitivity.

We demonstrate that rigidity homeostasis can be maintained across distinct initial conditions and, strikingly, that networks can even recover from severely disrupted states. Furthermore, we find that rigidity persists for a considerable range of disassembly rates and mechanosensitive thresholds, up to some critical values. Finally, we show that our models are close to the rigidity critical point for undercoordinated prestrained networks. The key steady-state topological and mechanical properties of the networks resulting from our sensitivity analysis are listed in Table II.

### A. Effect of initial states

To investigate if our models described in Sec. II can achieve rigidity homeostasis across distinct initial states, we use the steady state snapshot for the edge model (ESS) [Fig. 2(a)] as the initial state for the node model. Conversely, we use the steady state snapshot for the node model (NSS) [Fig. 5(a)] as the initial state for the edge model. All simulation parameters are kept the same as those in Table I. In both cases, the networks reach rigid steady states with exponential edge or node turnover at large *t* (SI, Fig. S1).

We find that the topological and mechanical properties depends on the number of the persistent elements, i.e., the number of nodes in the edge model, and vice versa. When ESS is used as the initial state and evolved to a new steady state under the node model, the time-averaged bulk and shear moduli and elastic energy are larger than the respective moduli and energy from the jamming initial state (Table II, Cases 2 and 4). The higher elastic moduli from ESS initial state is expected, as ESS contains more edges (*N*_*E*_ = 2714) than the jamming state (*N*_*E*_ = 2030), resulting in a denser network. On the other hand, when NSS is used as the initial state and evolved to a new steady state under the edge model, the resulting moduli and energy are lower than the respective values starting from the jamming initial state (Table II, Cases 1 and 3). The weaker elasticity attained via NSS can be explained by the fact that NSS contains fewer nodes (*N*_*V*_ = 1048) than the jamming state (*N*_*V*_ = 1359). At similar 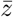, fewer nodes corresponds to a sparser network with smaller elastic moduli.

Because the nodes and edges in our models are diffusive, the remodeling dynamics is ergodic. Thus, it is expected that the same ensemble of network configurations is achieved at the steady state regardless of the initial configuration, as long as the node density is fixed in the edge model and the edge density is fixed in the node model. To verify this intuition, we run the edge model from yet another initial state: Based on a diamond network with *N*_*V*_ = 1000, *N*_*E*_ = 2000, 359 additional nodes are added at midpoints of random edges such that *N*_*V*_ = 1359, matching that for the jamming-derived initial condition. The evolution of the network reaches a steady state after 25,000 iterations. As shown by Cases 1 and 7 in Table II, the steady-state topological and mechanical properties closely agree regardless of the initial states.

The robustness of the steady state to initial configurations implies that our network models should be able to recover from drastic configurational changes. To test this hypothesis, we use a severely disrupted configuration with a large hole as the initial state for our node model. Starting from the NSS configuration [7(a, upper right)], we create a hole by splitting all nodes that lie in a spherical region of radius *R*_*H*_ centered at the origin, until only isolated edges remain in the hole. We set *R*_*H*_ = 2.4*ℓ*, which is 25% of the side length of the simulation box. Fig. 7(a, left) shows the snapshot of this initial state: Note the tensionless edges (white) in the hole (cyan) around the corners of the simulation box subject to periodic boundary conditions. As a result of this damage, the bulk and shear moduli and the elastic energy are 23%, 58% and 69% lower than the respective *B, G* and *E* for the NSS.

Under our node model, *B, G, E* rapidly recover to their steady-state values within 2*τ*_*V*_ [Fig. 7(b)]. We also verified that the structural properties *S*(*k*), 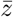 and *N* recover to steady states at similar time scales. A video showing the recovery process is given in the SI, Video S3 [59]. Fig. 7(a, bottom right) shows the network snapshot after 4.4*τ*_*V*_ (2000 iterations) starting from the damaged state. It is evident that the hole has been filled with tensioned edges and the structure is statistically indistinguishable from the NSS. These results are consistent with experimental and computational studies which suggest that catch-bonding crosslinkers facilitate the construction of stronger networks by mitigating crack initiation [38, 65].

**FIG. 7.**
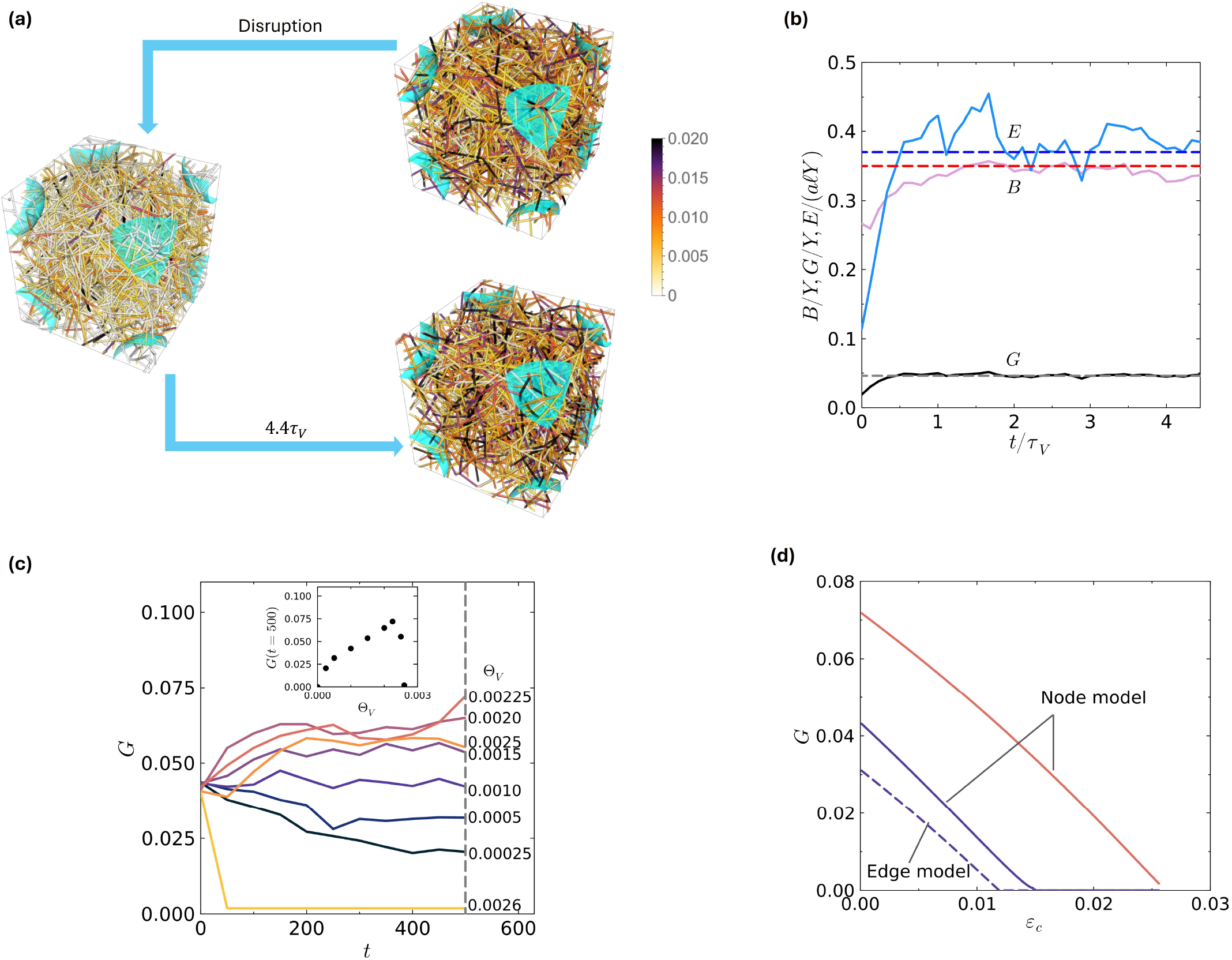
(a)–(b) Recovery of a damaged network under the mechanosensitive node model. (a) Snapshots of the damaged initial state with a large hole and the recovered network after 2000 iterations. The color bars indicate edge tensions. (b) Evolution of the bulk modulus *B*, the shear modulus *G*, and the elastic energy *E* starting from the initial state (a). The horizontal lines indicate the steady-state time-averaged *B, G*, and *E* (Table II, Case 2). (c) The evolution of shear modulus under different mechanosensitivity threshold Θ_*V*_, staring from the NSS [Fig. 5(a)] as the initial configuration. The inset shows *G*(*t* = 500) (vertical line) at different Θ_*V*_ . The network is stiffest when Θ_*V*_ = 0.00225. (d) Shear modulus of steady-state snapshots of the edge model with Θ_*V*_ = 0.001 (dashed, purple) and the node model with Θ_*V*_ = 0.001 (solid, purple) and Θ_*V*_ = 0.00225 (solid, red) upon compressional strain *ε*_*c*_ [Eq. (5)]. The networks are close to rigidity phase transitions.

### B. Effect of random disassembly rate

To study whether the rigidity homeostasis of our models is robust against variations in the random disassembly rate, we performed simulations of both models with *κ*_*E*_ = 1.5, 2 and *κ*_*V*_ = 0.75, 1, while keeping all other parameters in Table I unchanged.

As shown in Table II, specifically, by comparing Cases 1 and 5, and Cases 2 and 6, we find that in both models, *B, G* and *E* are robust against a remarkable 50% increase in the random disassembly rates *κ*_*E*_, *κ*_*V*_ . However, at even higher random disassembly rates *κ*_*E*_ = 2, *κ*_*V*_ = 1, we find that the tension- or force-sensitive disassembly processes begin to work against rigidity. Too many edges and nodes now fall below the thresholds Θ_*E*_, Θ_*V*_, such that the energy injection upon assembly cannot compensate for the loss of energy due to both random and tension-sensitive disassembly. A less stiff network results in more edges and nodes that fall below the thresholds. Such positive feedback eventually causes the networks to lose rigidity. The critical behavior of these rigid-to-floppy phase transitions is an interesting problem for future investigation.

### C. Effect of mechanosensitive disassembly threshold

The mechanosensitivity thresholds Θ_*E*_ and Θ_*V*_ determines which edges are disassembled due to their small tension. We have already seen in Sec. III that the networks lose rigidity if Θ_*E*_, *θ*_*V*_ = 0. Here, using the node model, we show that there is, in fact, an “optimum” value of Θ_*V*_ that builds the stiffest network given all other parameters fixed.

Starting from the NSS configuration [Fig. 5(a)] as the initial state, we run the node model with different values of Θ_*V*_ . Figure 7(c) shows the evolution of the shear modulus, and the inset shows *G*(*t* = 500) against Θ_*V*_ . The shear modulus initially increases with Θ_*V*_, and reaches a peak Θ_*V*_ = 0.00225, before dropping sharply. All simulations with Θ_*V*_ > 0.0026 lose shear rigidity in 10 iterations. This abrupt decrease in rigidity for Θ_*V*_ > 0.00225 is due to the aforementioned positive feedback: If too many edges fall below the mechanosensitive disassembly threshold, the network is expected to be less and less rigid in each iteration.

### D. “Distance” to rigidity critical point

The cytoskeleton is known to exhibit intermittent large fluctuations of stresses (“cytoquakes”), and it has been hypothesized that the cortex is at a state of self-organized criticality (SOC) that utilizes regulations of filament lengths and network connectivity to transition between subcritical and critical states in response to cellular needs [51, 66, 67]. In light of this hypothesis, we examine how close our networks are to the mechanical critical point at which tensional rigidity vanishes.

To determine the critical point, we start with the steady-state snapshots ESS and NSS and apply a compressional strain

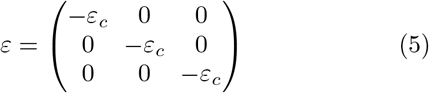

and monitor the shear modulus as a function of *ε*_*c*_, as shown by the purple curves in Fig. 7(d). The ESS and NSS models lose rigidity at 1.2% and 1.5% compression, respectively. We also performed the same analysis on the stiffest network found in Fig. 7(c), i.e., the node model with Θ_*V*_ = 0.00225, and found that even this network loses rigidity at 2.5% compression [Fig. 7(d), solid red curve]. These results demonstrate that networks with *z* ≈ 4 can emerge to be near criticality under mechanosensitive remodeling, consistent with the SOC picture.

## VI. DISCUSSIONS AND CONCLUSION

In this work, we study the actin cortex through the lens of functional metamaterials. The cortex in vivo is known to possess functions essential for the cell’s mechanical activities: maintaining a rigid steady state, recovering from structural disruption at arbitrary positions, and retaining a long-lived elastic memory. Achieving these functions together is challenging for synthetic polymer materials, and it is an unsolved puzzle how the actin cortex accomplishes them simultaneously.

By building two models that focus on assembly and disassembly of edges and nodes, respectively, we show that biologically-plausible mechanosensitive local rules lead to rigidity homeostasis, complete turnover, mechanical memory and near-criticality as collective emergent property of elastic networks. Specifically, one requires preferential disassembly that destroys low-tension elements, a small rate of random disassembly, and mechanisms associated with energy injection upon assembly. Together, these ingredients appear to be the minimum set that enable cortex-like properties, as summarized in Table III. Note that hyperstatic networks do not require prestressed filament to be rigid, but mechanosensitivity is still needed to achieve elastic memory.

**TABLE III.** Summary of models considered in this work.

| Model | Rigidity | Turnover | Elastic memory | Near criticality |
| --- | --- | --- | --- | --- |
| Hypostatic + Mechanosensitivity + Rand. disassembly | Y | Y | Y | Y |
| Hypostatic + Rand. disassembly | N | Y | N | N/A |
| Hypostatic + Mechanosensitivity | Y | N | Y | Y |
| Hyperstatic + Rand. disassembly | Y | Y | N | N |

A remarkable finding of our work is that nodes and edges undergo diffusion [Fig. 2(d), 5(d)] as demonstrated by the linearly increasing MSD with time well beyond the characteristic separation of nodes in the network. According to the Lindemann criterion, a solid should lose rigidity once the MSD of nodes exceeds the characteristic edge length. Clearly, the networks created by our models violate the Lindemann criterion egregiously–node displacements behave diffusively as if the system is fluid, yet it remains rigid with algebraic rheology, as observed experimentally [12, 14]. Our systems are no ordinary solids.

This observation, together with the fact that the nodes and edges undergo complete turnover, imply that despite steady nonzero elastic moduli, there are no permanently persistent correlations of the microscopic structure. This constantly shifting microscopic encoding (*i*.*e*., connectivity of nodes) of a macroscopic functionality (*i*.*e*., rigidity with stable elastic moduli) is known as “representational drift”, and has been found in numerous other contexts, including the visual and auditory cortices [68, 69], neural networks [70], and electrical Contrastive Local Learning Networks that learn AI tasks without a processor [71]. All of these systems share the common key property that they are abundantly *overparameterized* with far more tunable degrees of freedom than constraints. In our edge model, the tunable degrees of freedom are the binary ability of the edges to be there or not, while in the node model they are the binary ability of nodes to be there or not. For a network with *N*_*V*_ nodes and a mean valence of 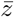, there are 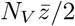 edges. The number of tunable degrees of freedom is therefore vast compared to the number of constraints (∼ 10^1^ components of the elasticity tensor) imposed by rigidity, namely, maintaining elastic constants within a narrow range.

This view of the actin cortex as tunable matter, with tunable degrees of freedom adjusted via rules arising from the mechanosensitive responses of various actin binding proteins, lies in stark contrast to the “use it or lose it” principle [5, 72] commonly used to understand the purpose of mechanosensitivity, *i*.*e*., that network constituents that contribute insignificantly to tension are selectively removed and redeployed so they might contribute more. In our view, Nature is not merely being thrifty–it is using local rules to tune degrees of freedom in order to achieve seemingly incompatible functions of rigidity homeostasis, structural healing, and elastic memory, which are important for survival.

One feature of tunable matter with local rules that adjust the tunable degrees of freedom, or “self-tunable matter,” is that the tuned functionality can be very robust. The ability to recover function despite partial damage is a hallmark of overparameterized systems that are tuned via local rules and that exhibit representational drift [45, 68, 71]. Very recently, it has been demonstrated that elastic networks equipped with mechanosensitive elements inspired by the actin cortex can be trained via a contrastive learning scheme [73]. Other types of mechanical networks as in vertex models of epithelial tissues have also been demonstrated to develop desired functionalities by adjusting tunable degrees of freedom using local rules [74–76]. In the future, we will extensively investigate how our models respond to different forms of internal and external stresses. An important next step is to combine our edge and node models to study if the elastic network achieves more robust rigidity homeostasis by incorporating multiple layers of local rules for adaptation and learning.

Our models provide prototypical platforms to build more realistic models that quantitatively match experimental data of the cytoskeleton, such as rheology and fluorescence recovery measurements. Since our goal here is to identify general “design principles” that enable rigidity homeostasis of networks under tensegrity, we have intentionally abstracted away molecular- and monomer-level details, including filament directionality and helicity, as well as the source of energy injection. We hypothesize that myosin motors and their interaction with treadmilling [77] could provide a valid mechanism to adjust the tensions of filaments and crosslinkers, adding additional tunable degrees of freedom that may increase robustness of rigidity homeostasis even further. When myosin is present, energy injection may not be coupled to assembly processes as we have assumed in our models.

We also remark that while the random disassembly processes are “hard-coded” into our models as the conceptually simplest mechanism to achieve complete turnover, cells living under fluctuating forces may not need to explicitly implement random disassembly, because tensions on filaments are already randomized, and it remains possible that mechanosensitive disassembly alone, together with assembly and energy injection, is another way to realize complete turnover. Furthermore, while we have successfully demonstrated that our models exhibit representational drift via diffusive dynamics of edges and nodes, the actin cortex are observed to have superdiffusive dynamics in certain conditions [14]. Finally, the form of stress decay in vivo is a low-exponent power law *t*^−*α*^ with *α* ≈ 0.25 [16, 17], which is even slower than the decay in our models; see Appendix B. Whether superdiffusion and small *α* arise from motor activities and the cortex’s adaptation to its fluctuating mechanical environment is an important direction for future research.

Our model assumes negligible bending rigidity, a choice justified by the cortical architecture. However, theoretical work suggests that a non-zero bending modulus blurs the sharp floppy-to-rigid phase boundary [51, 67, 78]. For a hypostatic network near 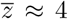, a nonzero bending stiffness ensures a floor in elastic moduli even at the stretching-controlled criticality. Rather than an abrupt failure at the critical point, this blurred boundary may provide the cell with a continuous mechanical control, allowing for highly tunable transitions between solid-like and fluid-like states.

New experiments can be designed to test several predictions of our models. One such possibility is observing rheology of cells immediately after a localized structural disruption, e.g., those occurring during endocytosis. Our “healing” simulations predict that the elastic moduli recovers within two characteristic turnover times. Another possible experiment is to measure the individual edge tensions using Föster Resonant Energy Transfer tension sensors [79] and determine the dependence of mean edge life time on their tensions. If the mechanosensitive disassembly is indeed a cornerstone of the cortex’s modeling, one expects to see that the edge life time is much shorter for low-tension edges.

Finally, we remark that our models are of fundamental interest for network theory, as the assembly and disassembly dynamics provide a novel mechanism to create evolving networks that undergo randomization of architecture while preserving network size, total edge length, and degree distribution, among other topological and geometrical characteristics. Compared to well-studied evolving networks that typically grow in size and usually become scale-free (i.e., power-law degree distribution) [80], it appears that many mechanical networks in living systems, constrained by energy and materials, favor homogeneity and more uniform distribution of node degrees. A fascinating avenue for future work is to study the evolutionary advantages of the topological features of cytoskeletal networks.

## DATA AVAILABILITY STATEMENT

Julia code for simulating both models and for characterization (e.g., linear response, structure factor, van Hove functions and mean-squared displacement), as well as all data used to produce in the figures and tables of this work are available on the GitHub repository Ref. [59].

## ACKNOWLEDGMENTS

We thank Emre Alca, Margaret Gardel, Paul Janmey, Makito Miyazaki, Michael Murrell, and Nathan Zimmerberg for instructive discussions. HW and MAGC were supported through the University of Pennsylvania Materials Research Science and Engineering Center (MRSEC), DMR-2309043, with additional support for HW provided by the University of Pennsylvania Center for Soft and Living Matter Postdoctoral Fellowship and for MAGC provided by the NSF through DMRMT-2005749. AJL acknowledges support of the Simons Foundation through Investigator grant #327939 and is also grateful for the hospitality of the Aspen Center for Physics (NSF grant PHY-2210452).

## APPENDIX

### A. Elastic moduli from linear response

To determine the elastic moduli of the network, we compute the stiffness tensor *C*_*ijkl*_ by measuring the variation of the stress tensor *σ*_*ij*_ under very small deformations (strains) *ϵ*_*kl*_ of the basis of the simulation box. The network is first relaxed to a local energy minimum. We then apply independent strains Δ*ϵ*_*kl*_ = 10^−5^ to each component of the basis. For every perturbation, the positions of nodes are re-optimized to allow for non-affine relaxations. The components of the elasticity tensor are obtained via finite differences:

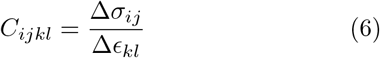

The resulting 9 × 9 tensor is mapped onto a 6 × 6 matrix using Voigt notations. We calculate the effective bulk modulus *B* and shear modulus *G* using the Voigt averages for a monoclinic material [81, 82]:

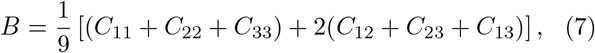

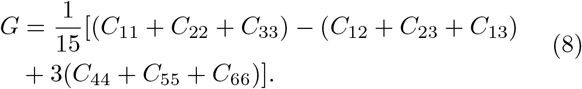

The equations for monoclinic materials are used because they apply to arbitrary simulation box basis vectors.

### B. Derivation of Relaxation Modulus from Microrheology

To extract the time-dependent stress relaxation modulus *G*(*t*) from microrheological data, we utilize the relationship between the dynamic shear modulus *G*^*\**^(*ω*) and the fluctuations of the network. We characterize the network dynamics through the Fourier transforms of the mean squared displacement of network components (nodes or edges) and stresses. For example, for the edge mode, where nodes are persistent, let 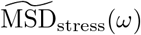 and 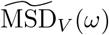 represent the MSD of stress and node displacements, respectively. The magnitude of the complex modulus is defined by the ratio:

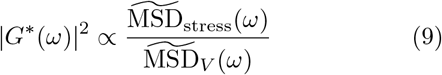

Because we operate in the quasistatic limit without viscous relaxation, the complex modulus *G*^*\**^(*ω*) is dominated by its elastic component. In this specific regime, the squared magnitude of the modulus in the frequency domain corresponds to the Fourier transform of the squared relaxation modulus:

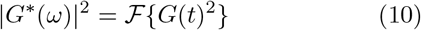

This identity allows us to recover the relaxation function *G*(*t*) by performing an inverse Fourier transform on the spectral ratio:

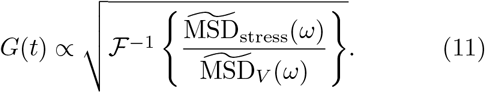

For both models, the stress MSD in our simulations approach a constant exponentially: MSD_stress_(*t*) ∝ 1 − exp(−*κt*), whereas the MSD for network components is diffusive. Applying these forms into Eq. (11) yields the large-time behavior

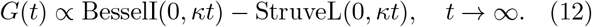

This function decays as *t*^−1^ at large *t*.

### C. Effect of steric repulsion

We have introduced steric repulsion in our models to compensate for the use of phantom edges. To investigate the effect of this treatment, we have also implemented control models without steric repulsion. Specifically, in our repulsion-free edge model, new nuclei are randomly generated uniformly in the simulation box. In our repulsion-free node model, orientations of dangling edges are chosen randomly.

Figure 8(a) shows the evolution of *B, G, E* of the repulsion-free node model, starting from the steady-state snapshot NSS [Fig. 5(a)]. Remarkably, these quantities remain positive and relatively stable for *t* up to 10*τ*_*V*_ . However, Fig. 8(b) shows that the structure factor at small wavevectors increase over iterations, indicating growing amplitudes of large-scale density fluctuations. We find similar density heterogeneity with the repulsion-free node model [Fig. 8(c)]. Thus, steric repulsion is not required for rigidity homeostasis with turnover, but is needed to maintain a *structural* steady state with density homogeneity.

**FIG. 8.**
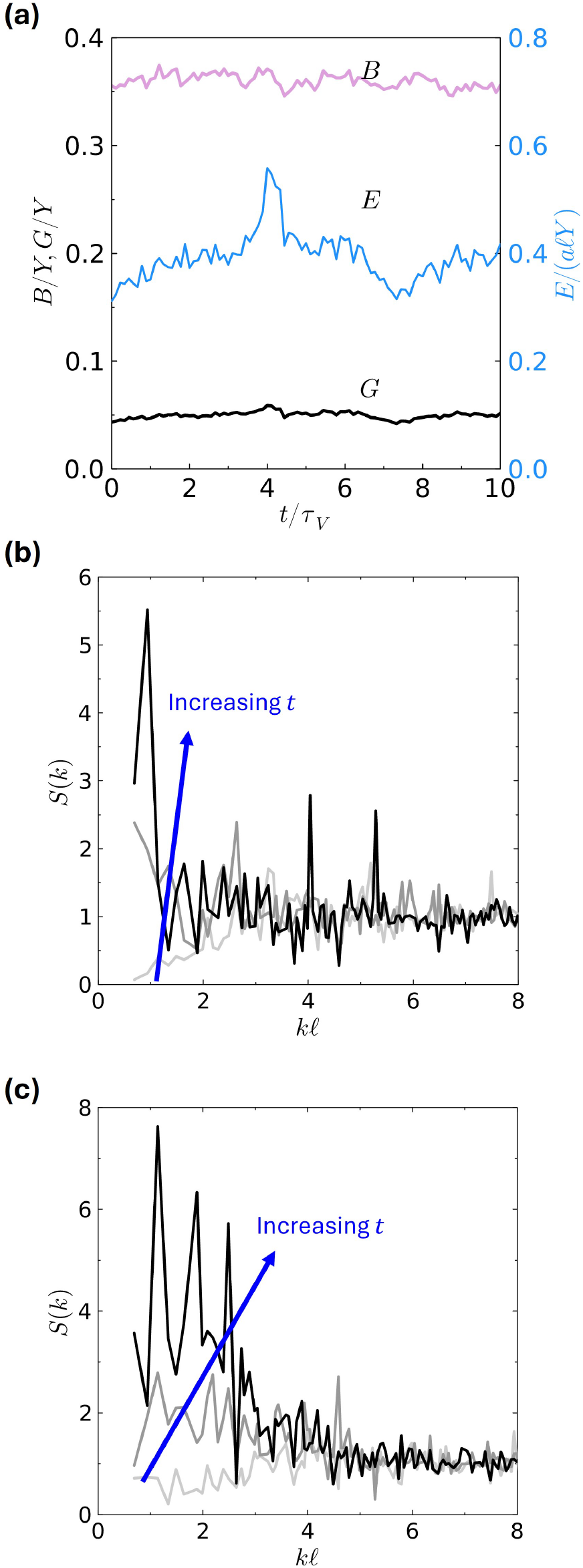
Effect of excluding steric repulsion. (a) Evolution of bulk modulus *B*, shear modulus *G* and elastic energy *E* for the repulsion-free node model, starting from the NSS configuration [Fig. 5(a)]. These quantities still reach a steady state. (b) Node structure factors for configurations at *t* = 0, 5*τ*_*V*_, and 10*τ*_*V*_ for the repulsion-free node model. (c) Edge-midpoint structure factors for configurations at *t* = 0, 5*τ*_*E*_, and 10*τ*_*E*_ for the repulsion-free edge model.

### D. Evolution of tension distribution

In the edge model, we observe the counterintuitive phenomenon that the elastic energy decreases while the bulk modulus *B* increases during the simulation. Since it has been shown that elastic networks with more homogeneous edge tensions tend to have higher *B* [61, 62], we investigate the evolution of tension distribution. Figure 9 shows the probability distribution function *p*(*ε*) of edge strains, which is equivalent to edge tensions measured in units of *aY*, at *t* = 50 and *t* = 10000. Indeed, we find that the edges have more uniform tensions as the network evolves.

**FIG. 9.**
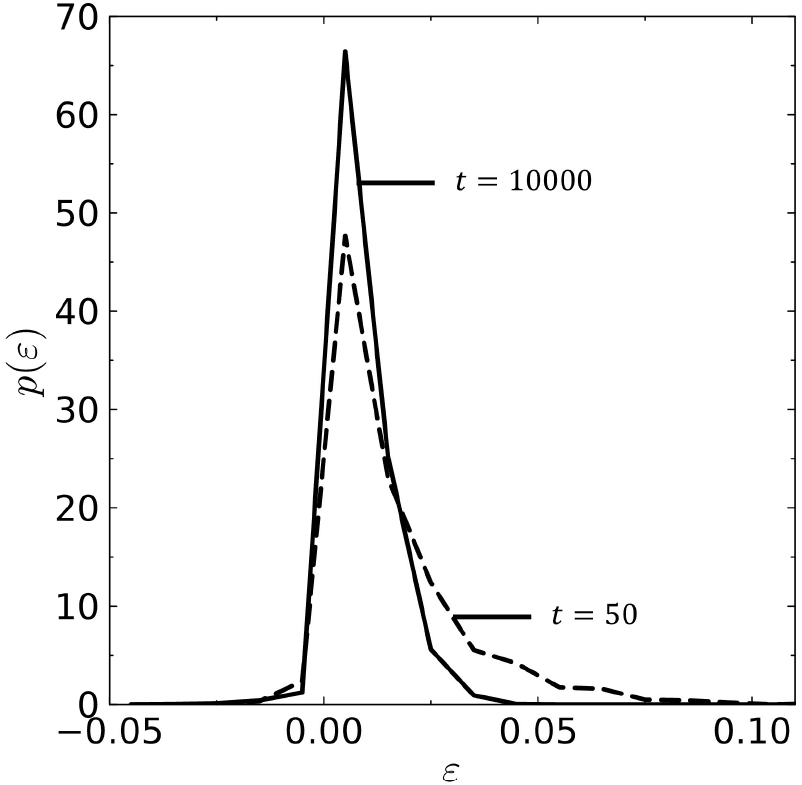
Probability distribution functions *p*(*ε*) of edge strains (tensions) at iterations *t* = 50 (dashed) and *t* = 10000 (solid) of the edge model, starting from the jamming initial state.

### E. Self van Hove correlation functions

The van Hove functions measure spatial and temporal correlations of evolving many-particle systems with *N* particles. The self van Hove function is the time-averaged probability distribution of particle displacements that take place during the time interval (*t, t* + Δ*t*). That is,

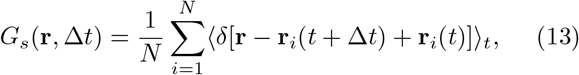

where **r**_*i*_(*t*) is the coordinates of particle *i* at time *t*. Figure 3 in the main text plots the van Hove functions for nodes in the edge model and edge midpoints in the node model, respectively. The distributions are spherically isotropic, so only displacements on the positive *x*axis are plotted. Both functions are heavy-tailed, and their forms can be accurately described by exponentialtruncated Lorentzian (red curves), given by

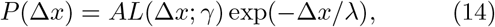

where *L*(Δ*x*; *γ*) = 1*/*[1 + (Δ*x/γ*)^2^] is the Lorentzian (Cauchy) distribution with scale parameter *γ, A* is a normalization constant, and *λ* is a truncation length scale. Figure 3 also plots the best-fit Gaussian and Lorentzian for the van Hove functions, from which it is clear that the simulated functions decay more slowly than Gaussians, but faster than untruncated Lorentzians. The heavytailed van Hove functions indicate that there exist a large fraction of anomalously fast and superdiffusive excursions of network constituents [15].

## Supplemental Information

### I. STATISTICAL TESTS FOR STEADY STATES

For the edge model, we performed two-sample t-test of *B, G, E* and *z* on two time windows *T*_1_ = [7001, 11006] and *T*_2_ = [14001, 18006], separated by 5.2*τ*_*E*_. We assume that two data points in each of these time series are considered independent if they are separated by at least *τ*_*V*_ . Thus, the effective sample size is the length of the time windows (4005) divided by *τ*_*E*_ = 1335, i.e., 3 independent samples per window.

For the node model, performed two-sample t-test of *B, G, E* and *z* on two time windows *T*_1_ = [7501, 9750] and *T*_2_ = [10001, 12250], separated by 5.5*τ*_*V*_ . The effective sample size is the length of the time windows (2250) divided by *τ*_*V*_ = 450, i.e., 5 independent samples per window.

For these tests, the null hypothesis is that the means in *T*_1_ and *T*_2_ are the same, and the alternative hypothesis is that the means are different. The results for *t*-tests are shown in Table S1. All *p* values are greater than 0.05, and thus the alternative hypothesis is not accepted in all tests, strongly suggesting that the test quantities reach steady states.

To further statistically validate that steady state is reached, we compute the symmetrized Kullback-Leibler (KL) divergence between the edge tension distributions in *T*_1_ and *T*_2_ :

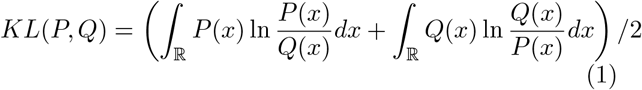

The KL divergence is 0.0099 for the edge model and 0.018 for the node model. For reference, we also generated two sets of uniformly distributed random variables on [0, 1], each with 2000 samples, close to the number of edges in our models. The KL divergence between these synthetic data sets is 0.014. Thus, the edge tension distributions between *T*_1_ and *T*_2_, separated by > 5 turnover times, are statistically indistinguishable.

### II. TURNOVER OF THE MODELS WITH DIFFERENT INITIAL STATES

Figure S1 shows the turnover of edges and nodes starting from different initial configurations. The turnover rate depends on the fraction of low-tension edges. For example, in the jamming-derived initial state, 0.07% of the edges have tensions less than Θ_*E*_ = 0.001. In the NSS, this fraction is 2%. Thus, in Fig. S1(a), the NSS has a much faster initial turnover in the edge model.

**Table S1.** Two-sample t-test for the edge and node models. The confidence intervals are for the difference in mean of quantities in *T*_1_ and *T*_2_.

| Edge model quantity | $p$ value | 95% Confidence interval |
| --- | --- | --- |
| $B$ | 0.16 | (-0.0023, 0.0098) |
| $G$ | 0.92 | (-0.0031, 0.0033) |
| $E$ | 0.99 | (-0.023, 0.023) |
| $z$ | 0.80 | (-0.0062, 0.0077) |
| Node model quantity | $p$ value | 95% Confidence interval |
| $B$ | 0.61 | (-0.012, 0.0073) |
| $G$ | 0.20 | (-0.0052, 0.0013) |
| $E$ | 0.08 | (-0.087, 0.0073) |
| $z$ | 0.73 | (-0.012, 0.017) |

**Fig. S1.**
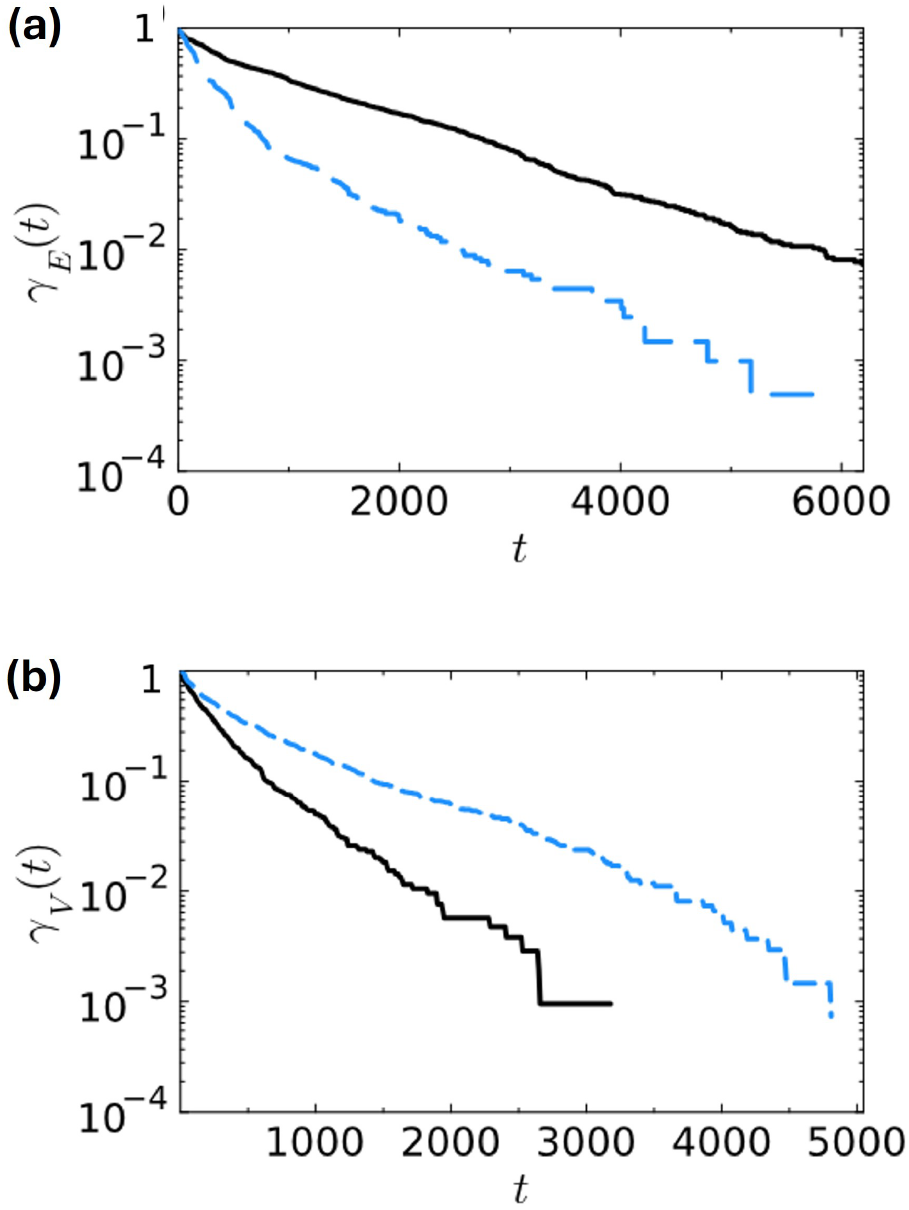
(a) Turnover of edges starting from the jamming-derived (black, solid) and node-steady-state (NSS) (blue, dashed) initial states. (b) Turnover of nodes for simulations starting from jamming-derived (black, solid) and edge-steadystate (ESS) (blue, dashed) initial states.

